# The vocal repertoire of white-nosed coatis: structural features, temporal dynamics, and social variations

**DOI:** 10.64898/2026.06.17.732959

**Authors:** Emily M. Grout, Christine C. Hass, Pranav Minasandra, Mara Thomas, Alison M. Ashbury, Josué Ortega, Margaret C. Crofoot, Ben T. Hirsch, Ariana Strandburg-Peshkin

## Abstract

For many group living species, vocal signals are a vital form of communication needed to coordinate social behaviours. Investigating these processes first requires a detailed description of the species’ vocal repertoire. White-nosed coatis (*Nasua narica*), which forage and move together in forest habitats, are thought to rely on vocalisations to coordinate movements and maintain group cohesion. However, quantitative studies of their vocal repertoire are lacking. We examined the vocal repertoire of white-nosed coatis from wild populations in Panama and Arizona, USA, to gain a more comprehensive view of their calling behaviours. By combining traditional acoustic analyses with an unsupervised approach based on spectrogram structure, we characterised the diversity of calls in this species and described the temporal and structural features of their vocalisations. We identified 19 call types, with some of these calls emitted in multi-syllable call sequences or in fast succession. We found variability in call rates among group members, which may be driven by differences in social status within the group. In addition, our results indicate that white-nosed coatis likely have individually-recognisable vocalisations. This study provides a foundational description of the white-nosed coati vocal repertoire, laying the groundwork for future research on vocal communication in this species.

## Introduction

Communication is fundamental to sociality, and can shape interactions that are critical for individual fitness. Vocal communication is particularly important for many group-living animals, as it allows individuals to rapidly share information across social contexts (Fichtel and Manser 2010). It also allows group members to flexibly adjust their behaviour in response to changes in their social and environmental surroundings. Vocal communication is especially valuable in visually restricted environments, such as dense forests, where vocal signals can help maintain group cohesion and coordinate behaviour when visual cues are limited (Teixeira da Cunha and Byrne 2008). By categorising the distinct call types used within a species, we can investigate how vocal signals function across different social and ecological contexts, providing insights into the evolutionary processes shaping vocal communication.

Establishing a species’ vocal repertoire has traditionally required prolonged observation from longitudinal studies of habituated animals (De Waal 1988; Gros-Louis et al. 2008; Salmi et al. 2013; Warrington et al. 2014). Yet, this approach is increasingly challenging in today’s research landscape due to the increased emphasis on novelty in publications and declining interest for long-term research among funding bodies (Lindenmayer et al. 2012). Recent advances in miniaturised audio recording technology are providing researchers with a new approach to collect large amounts of vocal data without requiring the habituation and long-term monitoring of populations (Couchoux et al. 2015). This is particularly useful for studying species in structurally complex environments where direct observation is limited.

Further, these technological advances provide new and exciting opportunities to examine long-term temporal dynamics of vocalisations across multiple individuals. By recording vocal sequences across members of social groups, researchers can investigate fine-scale call-and-response dynamics that were previously impossible through traditional observational methods. These continuous recordings also make it possible to uncover the behavioural contingencies underlying vocal communication, offering deeper insights into how individuals interact and respond to one another. Integrating these tools with traditional behavioural observations provides a framework for building a comprehensive description of a species’ vocal repertoire.

Call classifications have most commonly been made by visual inspection of spectrograms (Zimmermann 1985; Collias 1987; Warrington et al. 2014). Using these manual classifications, call features can be extracted from the spectrogram images (Collias, 1987; Crockford & Boesch, 2003). This method provides a detailed description of a vocal repertoire; however, visually classifying call types can be prone to human perception bias and spectrogram analysis is more limited in scalability and generalisability across large and diverse datasets. In recent years, unsupervised approaches have been increasingly adopted to distinguish call types (Dunlop, 2017; Thomas et al., 2022). These approaches reduce the need for labour-intensive manual annotation in large datasets, but may suffer from difficulty in interpretation. Despite there being limitations in both approaches, combining unsupervised approaches with manual classification can offer a more comprehensive and objective framework for characterising a species’ vocal repertoire.

White-nosed coatis are emerging as a valuable model system for studying within-group social and spatial dynamics (Gompper 1996, 1997; Hirsch and Gompper 2018; Grout et al. 2024).

They inhabit a diverse range of forested habitats across Central America, Mexico, and the southwestern United States and live in groups consisting of adult females and their offspring (Kaufmann 1962; Valenzuela and Ceballos 2000; Frey et al. 2013; Nigenda-Morales et al. 2019; Hass 2021). Groups spend the majority of their time foraging on the forest floor for fruit and insects, often splitting into smaller foraging parties composed of close relatives (Gompper 1994; McColgin et al. 2018; Grout et al. 2024). Adult males are typically solitary and are usually only found with groups during the mating season (Kaufmann 1962; but see Gompper and Krinsley 1992, Hirsch and Gompper 2017). Juveniles are the most aggressive group members, often monopolising food resources over sub-adults and adult group members (Kaufmann 1962; Hirsch 2007). Relatedness structures vary between groups, and there are often higher levels of within-group aggression between unrelated individuals (Gompper et al. 1997; Hirsch et al. 2012; Grout et al. 2024). Because they often inhabit forested environments and exhibit both differentiated social relationships and flexible grouping dynamics, coatis provide an ideal study system for examining how animals use sensory information to manage changing social and ecological conditions.

Despite being diurnal, coatis rely heavily on their olfactory and auditory perception, which may be a consequence of their nocturnal ancestry (Kaufmann 1962; Koepfli et al. 2007; Nigenda-Morales et al. 2019). Given that coatis often inhabit visually restricted environments, vocal communication likely plays a central role in their social dynamics.

Previous studies have shown that coatis produce a wide range of vocalisations, which are likely important in mediating collective processes such as coordinating movement, maintaining group cohesion, and responding to threats (Kaufmann 1962; Gilbert 1973; Smith 1977; Trudgian 1995; Maurello et al. 2000; Compton et al. 2001; Gasco et al. 2019; Hass 2021). While a detailed analysis on the acoustic properties of their close relatives, ring-tailed coatis (*Nasua nasua*), has been done (Gasco et al. 2019), information for white-nosed coatis is limited and contradictory.

The original Kaufmann (1962) study on coati behaviour and ecology describes roughly a dozen calls and the context in which they are used. Unfortunately, these observations were not accompanied by recordings or quantitative descriptions of the calls. It is, thus, difficult to match some calls to their original description. Studies that quantitatively described white-nosed coati vocalisations have focused on one or two call types in captive populations (Maurello et al. 2000; Compton et al. 2001). The *chirp* is the most well described call type (Gilbert 1973; Smith 1977; Maurello et al. 2000; Compton et al. 2001; Hass 2021). It is a contact call, grading in intensity, that is emitted when moving, foraging, socialising, and when excited (Maurello et al. 2000; Compton et al. 2001; Hass 2021). Although Kaufmann did not explicitly describe a *chirp*, his description of a *grunt* appears to match this vocalisation’s context. Four aggression calls have been described: *chitters*, *squawks*, *squeals,* and *growls. Chitters* are often emitted in rapid succession and as they increase in intensity, grade into *squeals* (Kaufmann 1962; Smith 1977; Hass 2021). The *squawk* has not been previously documented in wild populations, and since it was only observed in captivity, it is unclear whether this call is part of the natural vocal repertoire or a product of the captive environment (Compton et al. 2001). Several alarm calls have been described across studies. The *bark* was reported by Kaufmann (1962) and Hass (2021) as a short, loud call. *Grunts* have been described as a graded call occurring across contexts, ranging from contact to alarm situations, by Kaufmann (1962), Smith (1977), and Gilbert (1973). Variation in call terminology, along with the limited availability of recordings from wild populations, highlight the need for a comprehensive vocal repertoire analysis in natural settings to better understand the role of vocal communication in coati social behaviour.

While understanding a species’ vocal repertoire is key to uncovering how animals communicate, it is also important to determine whether individuals are able to recognise one another from their calls. Determining whether individuals have distinct vocal signatures has been a focus of many studies on vocal communication in social groups. The ability to distinguish group members can be critical for making the right decisions for who to associate with and who to avoid (Tibbetts and Dale 2007). In groups with differentiated social relationships, such as coatis, the ability to recognise conspecifics is especially important for avoiding aggressive individuals and providing coalitionary support to close social partners, such as relatives (Kaufmann 1962; Gompper et al. 1997; Hirsch et al. 2012). The ability to record all group members in the wild without the presence of human observers now opens new opportunities to investigate individual variation in vocalisations under fully natural conditions. A key challenge, however, lies in distinguishing whether acoustic differences reflect true individual signatures or artefacts introduced by the recording devices themselves. Consequently, it is essential to test and account for potential device-related effects when evaluating individual distinctiveness.

Here, we describe the vocal repertoire of wild white-nosed coatis, investigate temporal aspects of their vocal behaviour, and evaluate variation between individuals. We used a combination of audio logger recorders mounted on collars, manual hand-held microphones, and video cameras to collect recordings of vocalisations in two wild populations of white-nosed coatis in Panama and Arizona, USA. Handheld microphones were used to capture high-quality recordings for detailed descriptions of call types, while collar-based audio loggers recorded continuous data on vocalisations from group members in their natural habitat, providing unbiased sampling across time. To reliably classify call types, we integrated manual classification and unsupervised approaches. In doing so, we provide a comprehensive description of the white-nosed coati vocal repertoire which will facilitate the investigation into how vocal communication contributes to the social dynamics and collective behaviours of this species.

## Methods

### Study sites and data collection

Audio data were collected from wild white-nosed coatis in Panama and Arizona, USA, using a combination of recording methods. In Panama, small audio loggers (Soroka 18E, TS-Market; Edic Mini Tini+ A77, TS Market) were mounted to GPS collars (e-Obs Digital Telemetry, Gruenwald, Germany) worn by almost all members of one group (Galaxy) in Soberania National Park (SNP), and two adult females in one group (Smarties) on Barro Colorado Island (BCI). Methods for the capture and collaring protocol are in Grout et al. (2024). Coatis were categorised into four distinct age classes. Adults were defined as individuals older than 2 years. Subadults were those aged between 1 and 2 years. Juveniles were classified as individuals aged between 5 weeks and 1 year, and nestlings were defined as individuals younger than 5 weeks. The Galaxy group were composed of seven adult females, one subadult female, two subadult males, one juvenile, and one adult male (Grout et al. 2024). Audio data were collected from all members of the Galaxy group from December 25^th^ 2021 – January 6^th^ 2022 with a sampling rate of 24 kHz. The Galaxy group collars were equipped with two recording devices to extend the recording period. By combining data from both recorders, the audio loggers captured recordings from 0600 – 0900 for 14 ± 1.5 days. In February 2020, the two adults from the Smarties group wore collars each equipped with a single recorder, which sampled audio data at 22,050 Hz. These audio loggers collected data from 0600 – 1000 for 5 and 7 days. Sampling rates of the audio logger recorders mounted on collars were chosen to maximise the duration of the recording period while capturing a relevant frequency range. In February 2022, we recorded 20 minutes of audio data from a different wild group of seven coatis (Trago) in SNP. The Trago group were composed of one subadult male, five juvenile females, and one juvenile male. Recordings were collected using a hand-held ultrasonic microphone, at a sampling rate of 250 kHz. We also recorded audio data opportunistically when the Galaxy group members were trapped before we collared them.

Audio data were also extracted from videotape recordings of > 25 individuals across four groups of wild white-nosed coatis in Arizona, USA, between 1994 and 1999 (sampling rate 32 kHz), as well as audio collected separately as part of a video program (sampling rate 10 kHz; Sutor, 2000). Table S1 provides further details for each data stream.

Although the complete white-nosed coati vocal repertoire has not previously been described, past research has defined some of their call types. To maintain consistency with existing research, we categorised call types based on previous descriptions. For calls that were categorised differently between studies, we used the original names given. For calls that showed no similarity in acoustic structure to previously described calls, as determined by visual inspection of their spectrograms, we assigned a new classification. In this study, we manually extracted 61830 calls from the audio logger recordings that were mounted to the collars, 61657 of which were from the Galaxy group members (Figure S1 provides information on hours labelled per group member). From the manual recordings, we extracted 272 calls, with 179 from the Panamanian population and 93 from the Arizonan population.

### Ethical Note

Methods followed the ethical guidelines set by the American Society of Mammologists. Capture and collaring procedures conducted on Barro Colorado Island and Gamboa were performed in accordance with the Institutional Animal Care and Use Committees guidelines (Smithsonian ACUC clearance number: 2017-0815-2020).

### Analysis

#### Call feature extraction

To extract the structural features of each call type, one researcher (EG) labelled the start and end of a subsample of each call type by visualising the spectrograms in Adobe Audition (Adobe Inc. 2024). Spectrograms were generated using a 512-point fast Fourier transform (FFT) with a Hamming window. We selected the highest quality calls where there was no overlap of calls and the sample rate of the recording captured the full spectrum of the call.

Using the seewave (Sueur 2022) and tuneR (Ligges et al. 2013) packages in RStudio Version 4.0.3 (R Core Team 2017), we extracted the duration, dominant frequency, spectral flatness measure, as well as the interquartile range for the frequency distribution of each call. Calls which contained multiple components were split into each call component for acoustic feature extraction. To determine the variation of the structural features within each call type, we extracted a subset of call types (see Table S1 for sample sizes used). Calls were chosen based on their clarity, with priority given to those with minimal background noise to ensure accurate feature representation. Manual audio recordings were sampled at higher frequency ranges and exhibited lower background noise compared to audio logger recordings. Consequently, calls recorded from manual recordings were prioritised for spectrogram analysis. However, if certain call types were exclusively found in the audio logger recordings, their spectrograms were also analysed.

#### Manual call classification and unsupervised analysis

Calls from audio logger recordings from all collared group members in the Galaxy group were manually labelled. After filtering for call types which had more than 50 examples, we had 11 call types for mapping which totalled to 40816 calls (see Table S2 for all call counts). *Chirp grunt* and *chirp click*, which were frequently emitted two-syllable call sequences, were treated as unique call types in this analysis because these sequences potentially contained additional information compared to the call types, *chirp*, *grunt*, and *click*, treated separately. Calls were filtered to those whose durations were within the range of 0.03 to 0.8 seconds for realistic comparison (Figure S2).

As an alternative approach to human categorisation, and to test the robustness of our call classifications, we used an unsupervised clustering approach to visualise and assess the vocal repertoire based directly on call spectrograms, following the practical guide by Thomas et al. (2022). After generating spectrograms for each call, we converted these spectrograms into numeric vectors based on the intensity of sound in each time-frequency bin, then measured the distance between each pair of vectors as a metric of call dissimilarity (we refer to this as the *spectrogram distance*). To visualise the results, we then used the Uniform Manifold Approximation and Projection (UMAP) algorithm which uses these distances to map each call into a low dimensional latent space (McInnes et al. 2020; Sainburg et al. 2020), while preserving the relation of each spectrogram to others similar to it (i.e. keeping similar spectrograms close to one another in latent space). See supplements for more details on the spectrogram generation.

To assess the acoustic similarity between manually assigned call types, we used spectrogram-based distances in the original feature space (prior to dimensionality reduction). For each call, we identified its five nearest neighbours and recorded their call types. We then calculated, for each call type, the probability of each other call type being among its five nearest neighbours. These probability scores were normalised and log^2^ transformed to account for the variation in the frequency with which each call was emitted. As the nearest neighbour metrics only capture one measure of class separation in space, we also evaluated the distribution of all pairwise spectrogram distances within each call type compared to the distribution of pairwise spectrogram distances between different call types.

To visualise the acoustic similarity between different call types, we constructed a neighbourhood graph following the guidelines of Thomas et al. (2022). To construct the network, we transformed the evaluation matrix of the normalised five nearest neighbour probabilities from the original space embedding into a symmetric distance matrix. Each field was replaced with the average of itself and its diagonal counterpart (M[i,j] = mean(M[i,j], M[j,i]), and then M[j,i] = M[i,j]). The matrix was then multiplied by -1, and the diagonal was set to zero. Finally, we generated a graph approximation where edge length represented the distance values from the matrix, whereby larger distances represent greater acoustic dissimilarity between calls.

#### Temporal patterns in call sequences

We analysed inter-call intervals to assess the temporal patterns of white-nosed coati calling behaviour. This analysis only included calls from audio logger recordings of Galaxy group, as these recordings provided long, continuous sampling periods and captured a high volume of vocalisations suitable for detailed temporal analysis. To capture consistent patterns, we included only call types with 20 or more examples. Inter-call intervals were calculated as the time from the start of one call to the start of the subsequent call of the same type per individual, intervals were only used for cases where there were no other calls in-between.

We computed call transition matrices per individual by identifying all sequential call pairs that occurred within 20 seconds of one another. This 20-second cut-off was selected based on the distribution of inter-call intervals, as it captured 90% of these intervals. For each individual, we summarised the frequency of all observed transitions between call types, normalised the rows of each matrix to represent transition probabilities, and then averaged the resulting matrices across individuals to generate a group-level mean transition matrix.

#### Individual variation in call features and call rate

To examine individual differences in call acoustic features and calling behaviour, we used the same unsupervised method as described above for visual inspection of call similarities. We assessed the individual identities of the five nearest neighbours for all call types in the original feature space, assessing how frequently calls from the same individual were more acoustically similar to each other than expected by chance, regardless of call type. Because each individual had two recorders in the Galaxy group, we determined to what extent the recording devices influenced our results by running the same analysis with recorder identity. We then assessed the influence of the recording device on the acoustic similarity of the three most common call labels: *chirp*, *chirp grunt*, and *chitter*. Specifically, we tested whether calls from the same individual recorded on different devices were more similar than calls of the same type recorded from different individuals on different devices. To do this, we randomly drew 100,000 pairs of calls from the audio logger recordings, ensuring that all pairs of individuals and recorders had an equal chance of being represented, and computed the distribution of distances.

To gain a better understanding of how calls were used by group members within social groups, we compared the calling rates for commonly emitted call types for each group member in the Galaxy group. All data for this analysis were from audio logger recordings. All individuals had calls labelled from 0800 to 0900 between December 25, 2021, and January 6, 2022.We calculated call rates by summing the total number of calls of each type made by each group member, then dividing by the total number of hours that individual had labelled data. To ensure comparability, we restricted rate calculations to time periods during which more than 90% of group members had labelled data.

## Results

### Characterising call types

We characterised 15 call types in the white-nosed coati vocal repertoire (Figure 1), 12 of which had previously been described by other researchers, and three of which (*bop*, *hum*, and *pew*) were assigned a new classification due to not having similarities with previously described call types. We also observed five call types which were emitted from nestlings younger than five weeks old (*purr*, *mew*, *whine*, *whistle*, and *chitters*; see Figure S3 for spectrograms). Except for *chitters*, all of these calls were only observed in nestlings’ vocalisations. The calls emitted by adults varied in frequency range, duration (Table S3), and rate of emission (Figure S4). We found four multi-syllable call sequences as well as six call types which were emitted with consistent inter-call intervals (see Table S4 for further information and Figure S5 for spectrograms). Information on call contexts, based on previous literature, is given in Table S5. We found that five of the 15 call types had harmonics (*chitter*, *pew*, *squeal*, *squeak*, and *vibrate*) and the most frequently emitted call type, the *chirp*, had a biphonic component. We did not observe ultrasonic components in any of the fundamental frequencies of white-nosed coati calls. However, using the ultrasound recorders, we observed that the *chitter* and *squeal* harmonics often reached ultrasonic frequencies, with the frequency range of these harmonics varying based on the intensity of the fundamental frequency.

**Figure 1.**
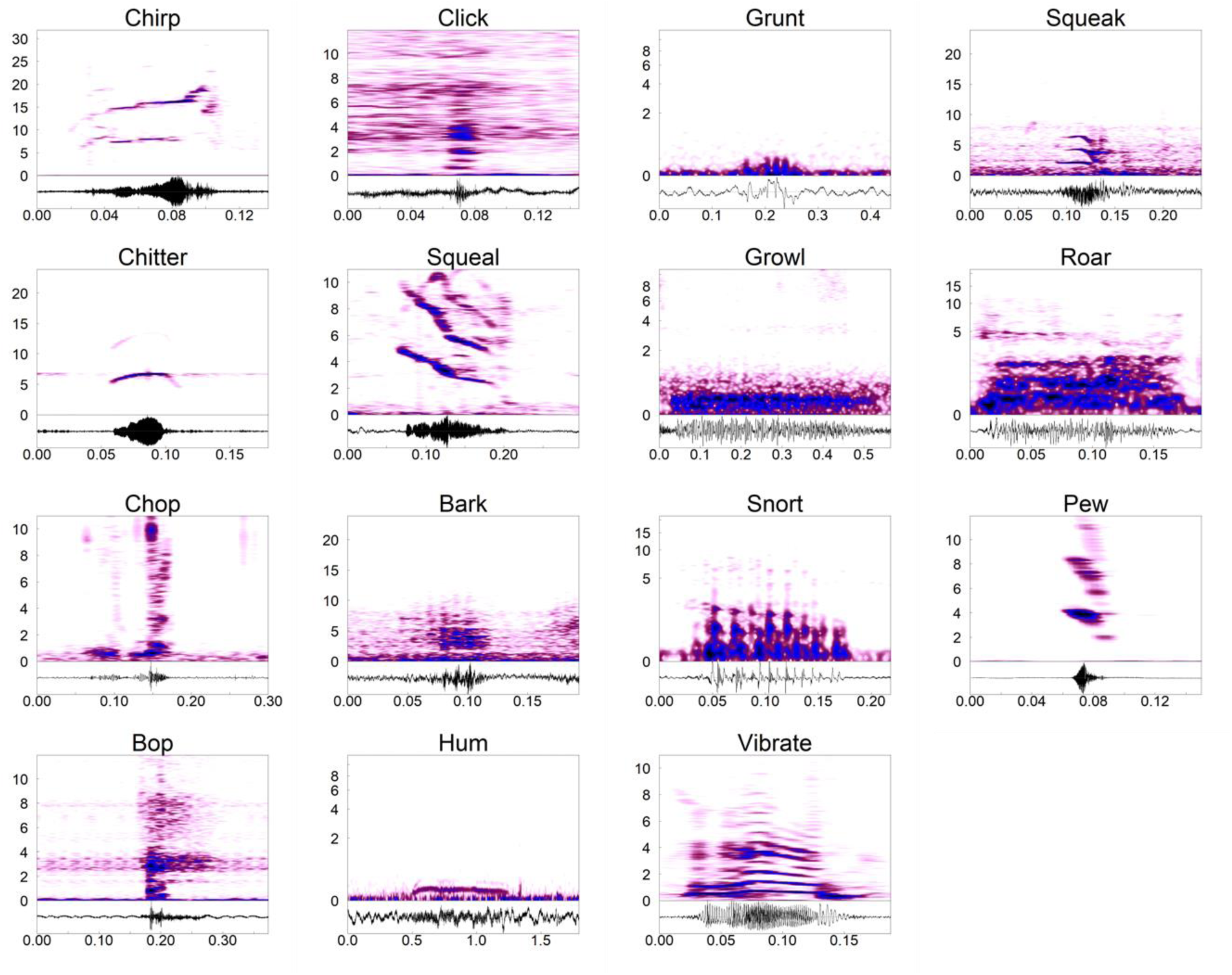
Spectrograms and corresponding waveforms of each call type emitted by adult white-nosed coatis. The *chirp*, *click*, *grunt*, *squeal*, *growl*, *roar*, *chop*, *snort*, *pew*, *bop*, *hum*, and *vibrate* call examples are from recordings from the Panama populations. The *squeak*, *chitter*, and *bark* call examples are from the Arizona population. For low frequency calls, the frequency scale (y-axis) is logarithmic.

### Unsupervised analysis of call types

Visually, the latent representation of calls showed distinct clustering of data into the observer-classified call types (Figure 2a). The *chirp* and *chirp grunt* had the greatest variation, while the *pew* and *chitter* had tighter clustering.

**Figure 2.**
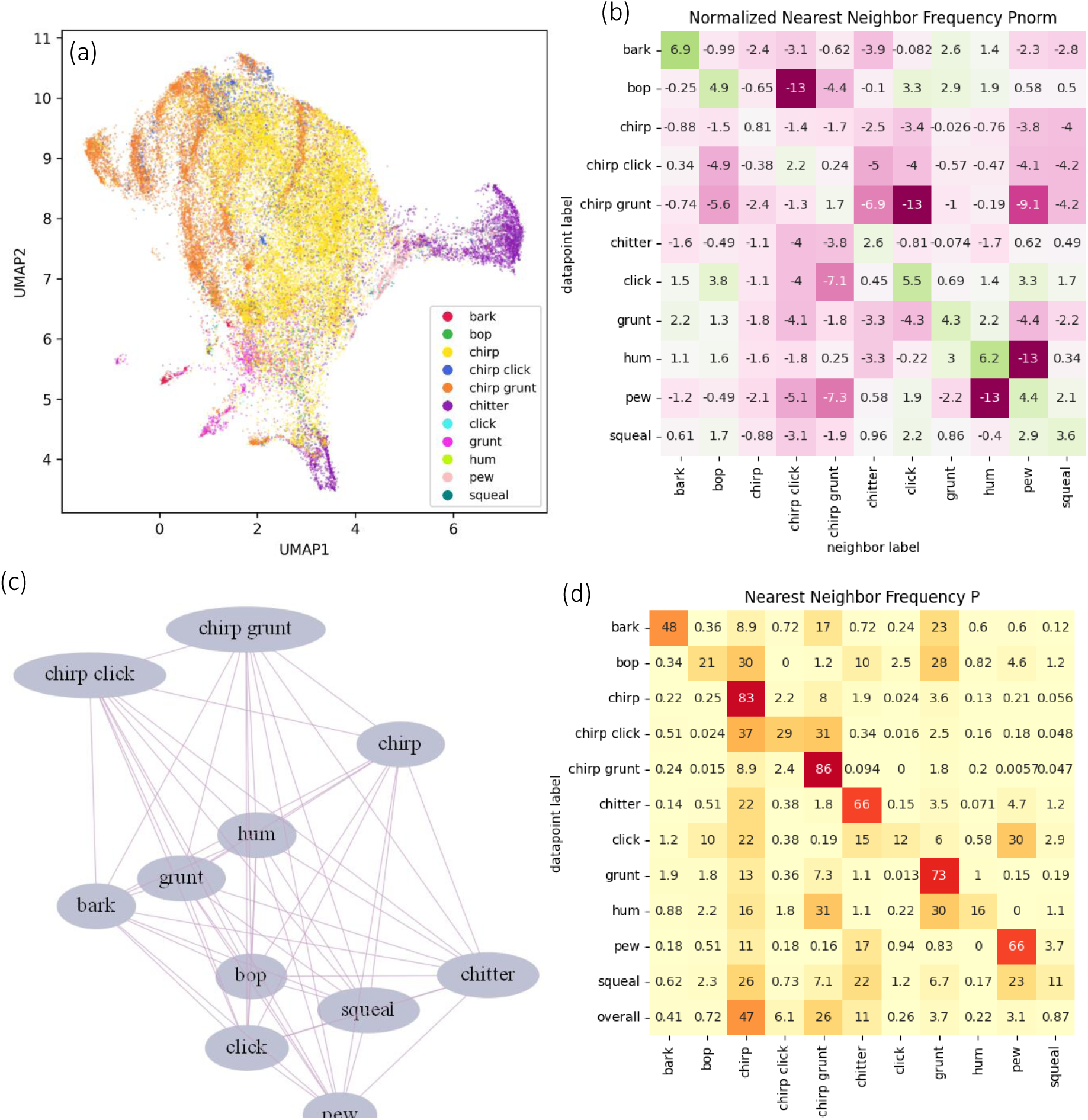
(a) Latent space representation of white-nosed coati vocalisations emitted by all members of the Galaxy group in Soberania National Park, Panama where datapoints are coloured by call types. (b) Matrix for the log2 transformed frequencies of each datapoint being within the five closest neighbours of each datapoint in original space. Values of two are four times as likely to be within the five nearest neighbours than expected by random chance, values of negative two are four times less likely to be within the five nearest neighbours than expected by random chance. (c) Neighbourhood graph where nodes represent call types and edge lengths represent the distance values from the log2 transformed matrix. (d) Matrix of the observed probabilities (given as percentages) of each call type being within the five closest neighbours of each other call type in original (before dimensionality reduction) space. The overall percentage of call types in the dataset is shown in the final row.

### Assessing call similarity with nearest neighbour metrics

The evaluation score, representing the unweighted average of the same-class probability for all classes in the original space, was 49.6, while the normalised evaluation score was 2.5. This means that across all data, for the five nearest neighbours of a given call, 49.6% of those calls were of the same type, and that calls of the same type were on average 2.5 times more likely to be within the five nearest neighbours of a given call than expected by chance. For the commonly emitted calls such as the *chirp, chirp click, chirp grunt, chitter,* and *squeal,* we observed little difference in pairwise distance between spectrograms in original space (Figure S6), but these differences were observed in the 2D UMAP space (Figure S7).

The percentage of same-class neighbours varied among different calls (Figure 2d). For instance, the *chirp* call had 83% same-class neighbours, though this call type represented 47% of the total dataset. When normalised and log2 transformed, its score was still positive (0.81), indicating that calls of this type were more likely to be neighbours than expected by chance. However, this score was less than for other call types such as the *bark*, which had a normalised log2 value of 6.9, making it 119 times more likely to be found near calls of the same type than expected by chance. When inspecting the normalised log2 transformed matrix, we found high similarity in some of the spectrograms of the call types (Figure 2b). The *bop* was a rare call type in our dataset (N = 293, <0.01% calls in our dataset), and was closely associated with the c*lick, grunt,* and *hum*. The *click* was closely associated with the *pew* and *bop*, and the *chitter* was more closely associated with the *squeal*. We found that the *chirp, chirp click,* and *chirp grunt* were distinct from one another in the normalised log2 transformed matrix, however the *chirp click* in the overall percentage of nearest neighbours was closer to the *chirp* and the *chirp grunt* compared to other *chirp clicks*.

The neighbourhood graph highlights acoustic similarities between call types (Figure 2c). It shows that the *chirp grunt* and *chirp click* are structurally closer together, likely because they both contain the *chirp* call. The network also indicates that the *chitter, squeal*, *click*, *bop*, and *pew* are structurally closer together. Additionally, the *bark* and the *grunt* are structurally similar to each other, which is likely due to their comparable fundamental frequencies.

### Call transitions and inter-call intervals

Of the 11 call types with sample sizes sufficient for analysing inter-call intervals and call transitions (>20 samples; Figure 3c), four (*squeal*, *pew*, *chitter*, and *bark*) exhibited smaller interquartile ranges for inter-call intervals, indicating greater temporal consistency in their emission (Figure 3a and 3b). We found that when one of these four call types was emitted, it was followed by the same call type in the majority of cases (83.4 ± 10 %) (Figure 3d). Of these four calls, we observed that the time intervals between *bark* calls exhibited the longest duration between successive calls (median = 0.27 ± 0.06 s). We found that the *pew* had the shortest inter-call interval and had the least variation in these durations (median = 0.11 ± 0.04 s). The *squeal* and *chitter* had slightly greater variation in inter-call intervals (0.16 ± 0.18 s, 0.16 ± 0.15 s respectively). The mean sequence length for each call type, where sequences contained no other call types, was 5.87 ± 1.66 s for chitter, 5.30 ± 1.89 s for *pew*, and 2.63 ± 1.16 s for *squeal*, with maximum call counts reaching 80, 47, and 39, respectively (Figure S8).

**Figure 3.**
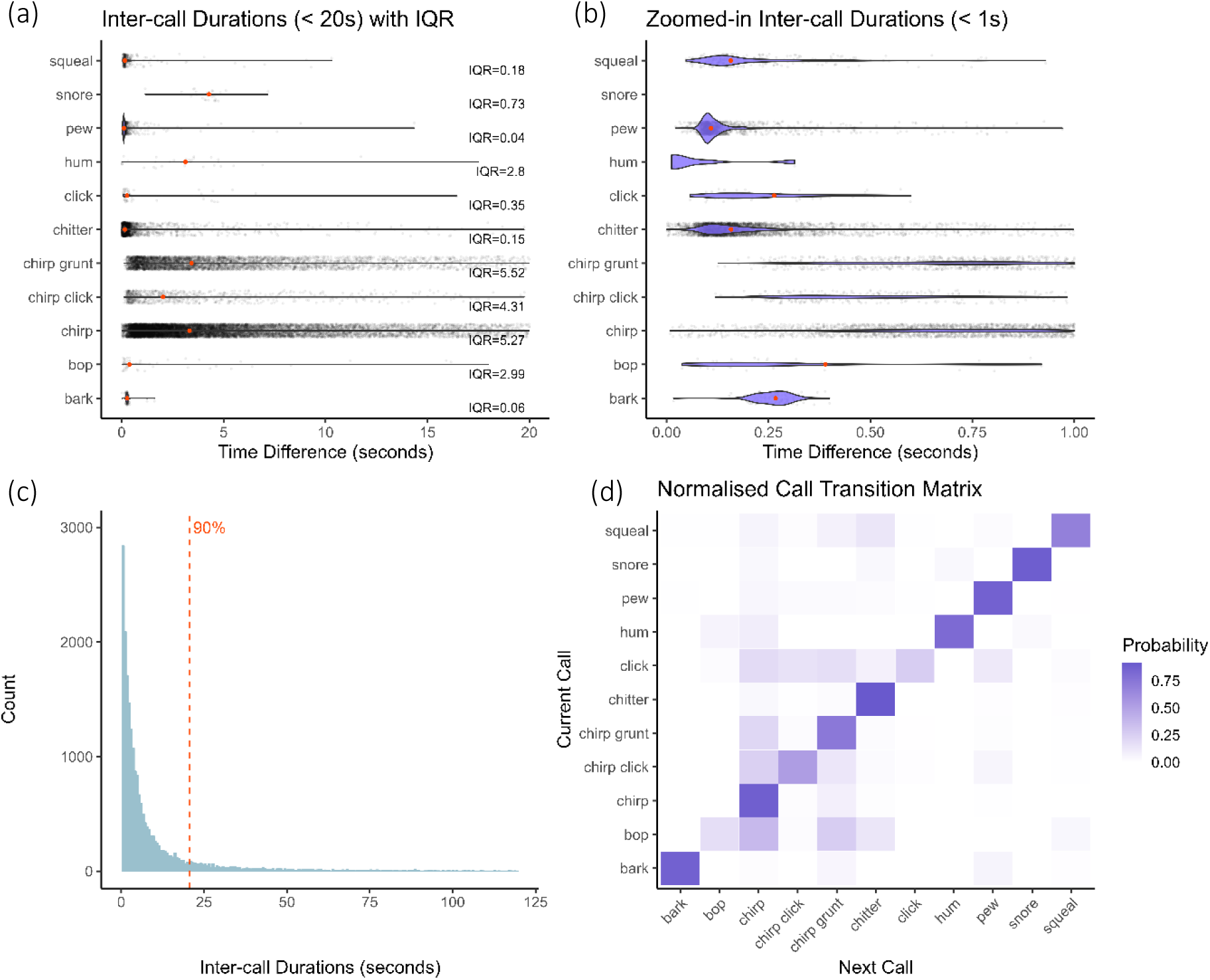
All plots show data from vocalisations recorded from all 11 collared white-nosed coatis in the Galaxy group, Soberania National Park, Panama. Only call types with 20 or more occurrences were included in these plots. (a) Distributions of within individual inter-call durations with the interquartile range. Raw data points are in black; red points indicate median durations. (b) Expanded 1-second segment from panel (a). (c) Histogram of all inter-call durations. Dashed red line indicates the 90th percentile threshold. (d) Normalised transition probability matrices for included call types.

### Individual variation

The unweighted average of the same-individual probability across all classes was 49.6%, with a normalised evaluation score of 2.465, suggesting that calls from the same individual tended to be more acoustically similar (i.e. shorter spectrogram distances) than those from different individuals (Figure 4a and 4c). Since each individual was fitted with a collar containing two audio recorders, we tested whether this similarity could be due to recorder-specific acoustic features, rather than vocal individuality. By repeating the analysis using the recorder ID as the grouping factor, we found that the same-recorder probability was 43.8%, with a higher normalised evaluation score of 3.307 (Figure 4b and 4d). As the normalised score exceeded that of individual identity, this suggests that acoustic similarity in the latent space is at least partly driven by artefacts introduced by the recording equipment rather than by individual vocal signatures. We found that the distribution of spectrogram distances for the *chirp*, *chirp grunt*, and *chitter* was slightly smaller within individuals recorded on different devices compared to between individuals, though the distributions largely overlapped (Figure S9).

**Figure 4.**
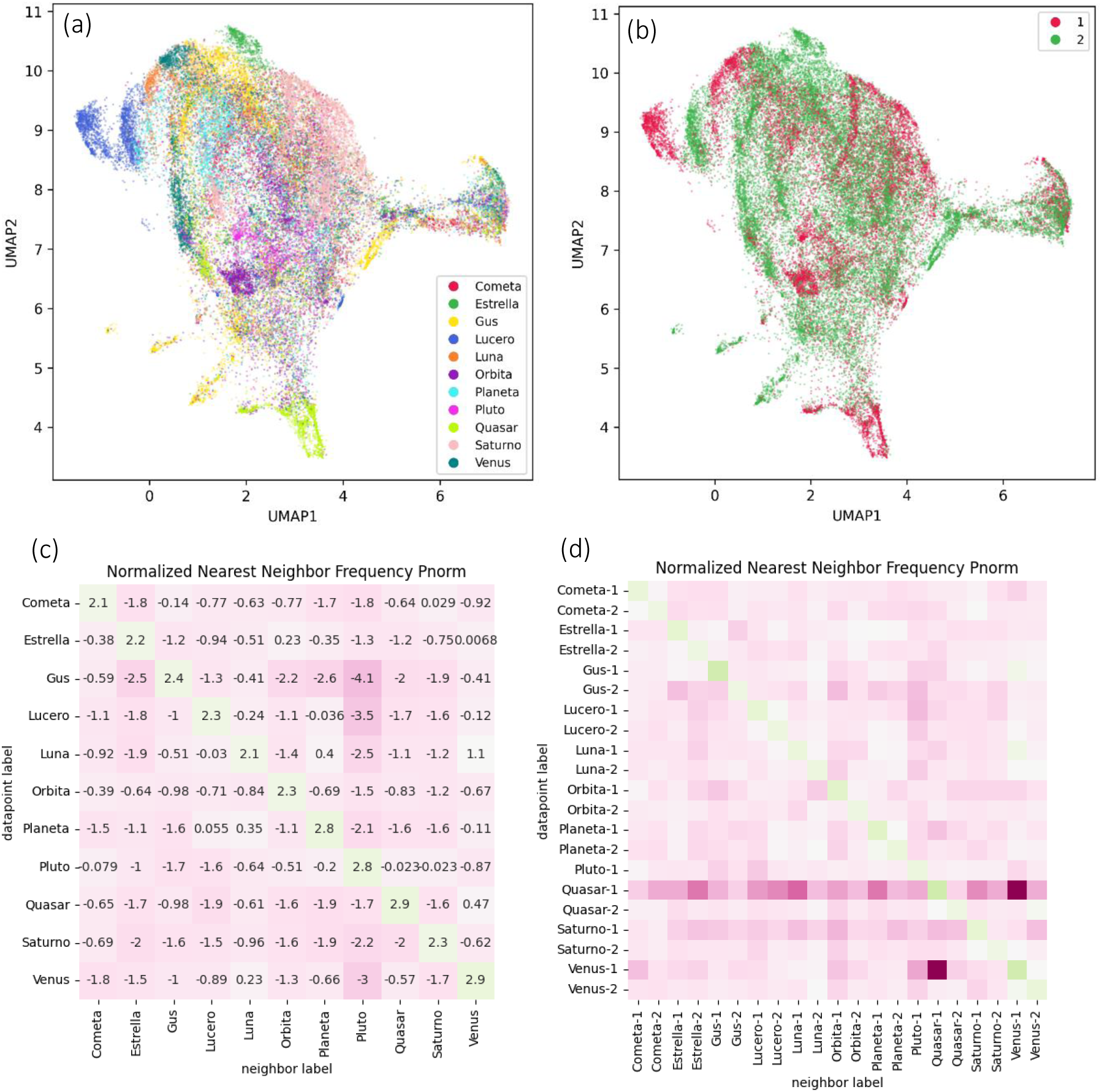
(a) Latent space representation of the most commonly emitted white-nosed coati vocalisations from all 11 collared members of the Galaxy group, Soberania National Park, Panama where data points are coloured by individual identity. (b) The same latent space representation as in (a), but with data points coloured by recorder identity for each group member. (c) Matrix for the log2 transformed frequencies of each group member’s datapoint being within the five closest neighbours of each datapoint in original space, regardless of call type. Values of two are four times as likely to be within the five nearest neighbours than expected by random chance, values of negative two are four times less likely to be within the five nearest neighbours than expected by random chance. (d) The same analysis as in (c), but with results separated by recorder identity for each individual.

We observed variation in the calling rates for each call type between different age-sex classes (Figure 5). Among the adult females, there was variation in the call rates of *chirp* and the *chirp grunt* calls (see Figure S10 for individual differences). The adult male emitted the *bark* and the *pew at* the highest rate among group members. The juvenile female had the overall lowest calling rates but emitted the *chitter* call most frequently amongst the group members. One of the subadult males had the highest calling rate of all group members, on average emitting the *chirp* call more than three times per minute. The *bop*, *grunt, hum,* and *click*, were emitted least frequently for all group members.

**Figure 5.**
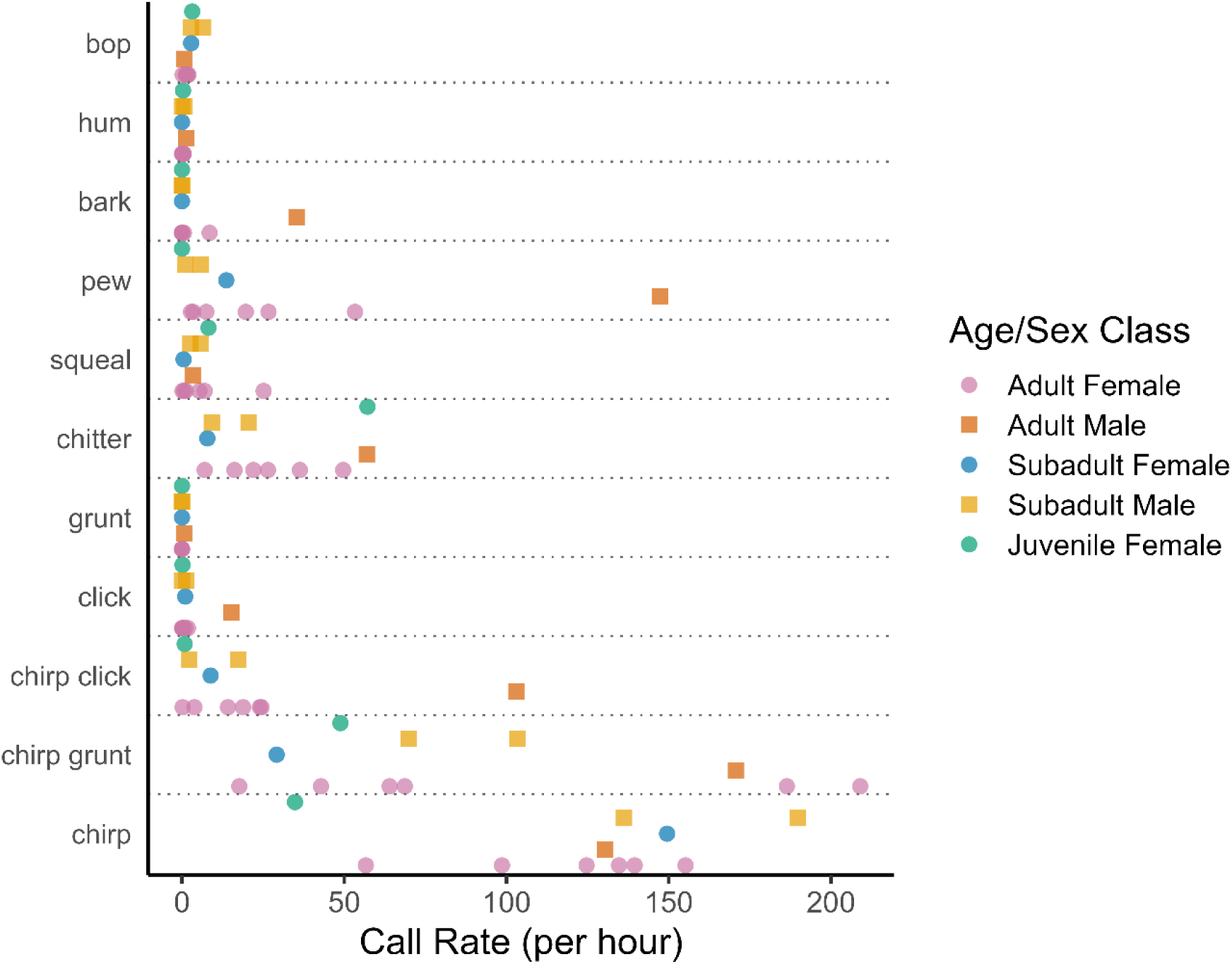
Call rates of all 11 collared white-nosed coatis in the Galaxy group in Soberania National Park, Panama. Each point represents the mean call rate for each individual for one call type.

## Discussion

Our manual classification of wild white-nosed coati vocalisations identified 19 distinct call types, with 15 predominantly emitted by adults, subadults, and juveniles, and four only produced by nestlings. We found that our manual classifications from audio logger recordings aligned well with an unsupervised analysis based on spectrogram distances, providing support for the reliability of our manually-defined call types and also highlighting the value of unsupervised approaches for validating human-defined call categories.

Furthermore, we showed that some calls (chitter and squeal) are emitted in fast succession with variable inter-call durations indicating that calling frequency may be modulated according to context and provide additional information to receivers. Although on-collar recorders accounted for much of the structural variation between individuals’ calls, we found marginal evidence for individual vocal signatures in the most commonly emitted call types. For the more commonly emitted call types, previous observations have provided evidence for their contexts, while rarer call types which have not previously been described require further investigation to determine their usage and significance.

Below we discuss the behavioural contexts associated with each call type, drawing on both previous reports and our own observations. We then address call types previously described but not observed in our study, as well as novel call types we have identified. This is followed by an examination of potential adaptations in the structural features of coati vocalisations in relation to external factors, and a discussion on individual differences in call rate emission and individuality encoded in call structures. Finally, we highlight the benefits of our methodological approach for describing a species’ vocal repertoire, and outline future research directions.

### Call Contexts

#### Contact Calls

There are multiple coati calls defined as a contact call based on their function. The *chirp*, *grunt*, *chirp grunt*, and *chirp*-*click*-*grunt* are emitted when foraging, moving, and resting (Kaufmann 1962; Trudgian 1995; Maurello et al. 2000; Compton et al. 2001; Gasco et al. 2019; Hass 2021). They can also be emitted when surprised, or when greeting others (Hass 2021). It is likely that the combination of these calls emitted and the rate of emission provide contextual information (Trudgian 1995; Hass 2021). Previous observations have suggested that the *click* and *grunt* components of the complex call are emitted when the group is moving or about to move (Hass 2021). These lower frequency call components can travel farther, making them more detectable by group members. Emitting these calls when initiating group movement is likely to increase their effectiveness in coordinating group members to leave. We found that there is variation in the calling rate, fundamental frequency, amplitude, and duration of the *chirp*, which may signal an individual’s motivational state. Previous examination on this call type has suggested that energy given to the different components varies with circumstance (Hass 2021). The multi-component structure of the *chirp* may allow flexibility in the information transferred.

#### Excitement call

One of the rarer call types in our dataset, observed to be emitted when an individual was excited, is the *squeak* (Hass 2021). The *squeak* is a tonal, relatively high frequency call. Although the *squeak* was rarely observed in the Panamanian populations which had audio logger recorders, we found that it was emitted by adult females in a wild population in Arizona. This suggests that there may be differences in call usage between study populations, and highlights the importance of sampling across a species’ range to obtain a more comprehensive understanding of the diversity and function of calls emitted within the species.

#### Alarm Calls

In line with previous studies, we identified different alarm calls which varied in intensity and circumstance (Kaufmann 1962; Gilbert 1973; Smith 1977; Gasco et al. 2019; Hass 2021).

The *bark*, a broad band alarm signal, is typically given when a coati is startled (Kaufmann 1962). This call serves to warn others of the potential threat, prompting group members to leap into the trees and become vigilant (Hass 2021). The individual closest to the source of the alarm often continues *barking* at the threat, while raising its nose and flicking its tail (Kaufmann 1962; Gilbert 1973; Smith 1977; Gasco et al. 2019; Hass 2021). We found that *barking* can have added elements such as the *chirp* before the *bark*, or the *grunt* during the *bark,* which may provide information on the caller as well as urgency. The *chop*-*chop* is emitted in fast succession by rapidly opening and closing the mouth repeatedly, previous studies found this call to only be emitted by adult males during the breeding season (Kaufmann 1962; Smith 1977), whereas in our observations it was also given during animal capture, suggesting it is produced more generally in threatening situations. Similarly, *snorts* and *roars* were only recorded when coatis were trapped, indicating these calls are likely associated with alarm, aggression, and distress.

#### Aggression Calls

Foraging competition is one of the main drivers for aggressive interactions in coati groups (Kaufmann 1962; Hirsch 2007; Hirsch et al. 2012; Hirsch and Gompper 2018). However, aggression can also arise from social pressures, such as competition between males for mating opportunities and conflicts over group membership (Hirsch and Maldonado 2011; Hirsch et al. 2012). The intensity of these agonistic interactions is shaped by several factors such as group spread, the number of individuals involved, the quantity and quality of the contested resource, and individual hunger levels. Given this variability, it is likely that coatis modulate their vocalisations in response to the nature and intensity of these conflicts, resulting in acoustic variation across aggressive contexts. *Chitters*, *squeals*, and *growls* have been described as aggression calls in previous studies (Kaufmann 1962; Smith 1977; Hass 2021). We found *chitters* to be the most frequently emitted aggression call, ranging from short single calls to loud series given in rapid succession. *Squeals*, which we found to be structurally similar to *chitters*, often accompany *chitters* during escalated fights (Kaufmann 1962). There was greater variation in the distribution of inter-call intervals for *chitter* and *squeal* calls further supporting our expectation that these aggression calls grade in correspondence with the severity of the aggressive interaction. Interestingly, we found that juvenile *chitters* clustered separately from adult *chitters*. This either suggests that juveniles use these calls in different contexts or that the structural differences are due to their not yet fully developed vocal apparatus. Future studies should include behavioural observations of *chitter* calls emitted across age groups to determine the cause of these structural differences.

We found variation in the acoustic properties of aggression calls. *Chitters* and *squeals* are tonal and high frequency, whereas *growls* are broadband and low frequency. This suggests that while all three call types are used in aggressive contexts, their differing acoustic properties may serve distinct functions. The motivation structural rules hypothesis suggests that high frequency, tonal calls should be used in friendly/fearful contexts, whereas low frequency, broadband calls should be used in aggressive contexts (Morton 1977). This suggests that high-frequency chitters and squeals may be used to signal distress or alert nearby group members, whereas low-frequency growls could be employed to intimidate opponents or assert dominance during aggressive interactions. Future studies should collect contextual information during aggressive interactions to test this hypothesis.

We found that *chitters* have harmonics reaching ultrasonic frequencies. Coatis can detect sounds up to 95kHz, with their most sensistive hearing up to 45 kHz (Peterson et al. 1969). Detecting these harmonics may be important for assessing the scale of within-group aggressive interactions, highlighting the need to sample vocalisations across a broad frequency range to accurately capture the nuances of social communication in this species.

#### Comparison with Previous Vocal Descriptions

The only call types we did not find in our recordings, which has been previously described in the white-nosed coati literature were *chuckling* and the *squawk* (Kaufmann 1962; Compton et al. 2001). The *squawk* was described as a loud, harsh call produced during agonistic interactions, characterised by a low frequency and relatively long duration (Compton et al. 2001). This call type has only been discovered in a captive population, therefore it is possible that this call represents a modified form of the *chitter*, *squeal*, or *growl*, which are emitted during agonistic interactions in wild populations. *Chuckling* is described as a series of repeated calls, including *chirps*, *chitters*, and *squeaks*, each emitted in brief sequences before switching to a new sound (Kaufmann 1962; Gilbert 1973; Hass 2021). It is most frequently emitted during the breeding season but also occurs throughout the year during coati reunions, mutual grooming sessions, and when captive coatis greet their handlers (Kaufmann 1962; Gilbert 1973; Hass 2021).

We identified three call types not characterised in previous studies which we have named the *bop*, *hum*, and *pew*. The *bop* and *hum* are soft, low-frequency calls that have only been detected by on-collar recorders. Their low amplitude may be the reason that these calls were not detected in previous studies that did not employ collars. The *pew*, observed only in the Panamanian population in this study, exhibited a structure distinct from other call types, closely resembling aggression calls based on our unsupervised analysis. This call was frequently emitted in rapid sequences, with variations in intensity and bout duration, indicating a potential link to the signaller’s excitement or motivation. We hypothesise that the *pew*, predominantly emitted by the adult male during the breeding season in the Galaxy group, serves a similar function to *chuckling* and may occur within the *chuckling* bouts observed in other populations. If these vocalisations are linked to heightened arousal, variation in call use during male–female interactions likely occur across individuals and populations. Further research, including behavioural observations, is necessary to fully understand the function and context of these newly classified call types.

#### Influences on Vocal Structures

We found that the most frequently emitted call types - *chirps*, *chirp grunts*, and *chitters* - have short durations and high fundamental frequencies. In forested habitats, high frequency calls rapidly attenuate, especially calls of shorter durations, therefore we hypothesise that these calls are an adaptation to minimise their detectability by predators, as observed in other species (Wilson and Hare 2004; Ramsier et al. 2012). Additionally, the frequency range of these calls (> 6kHz) are similar to many bird species inhabiting the same area, potentially providing further acoustic camouflage from predators (Hass 2021). The low-frequency calls, such as *growls* and *roars*, were rarely emitted, suggesting they are reserved for situations such as intimidating a competitor or a predator (Kaufmann 1962; Hass 2021). This relationship between call frequency and duration highlights the potential adaptive significance of vocal behaviour in white-nosed coatis, balancing effective communication with predation pressures.

#### Individual Variation

Our results on individual identity show that the recording device had a large influence on the similarities between calls, highlighting the importance of using the same recording device to test whether individuals have vocal signatures. Although recorder identity was a major confounding factor, we found that the most common call types—*chirp*, *chirp grunt*, and *chitter*—were marginally more similar within individuals recorded on different devices than between individuals. This is consistent with white-nosed coatis having individually distinct vocalisations, although further research is needed to determine whether they can recognise one another by their calls alone. Vocalisations recorded from captive populations found structural variation in *chirp* calls between individuals which further supports that they do have individual signatures (Maurello et al., 2000; Trudgian 1995). In a study of two captive populations using recorders which sampled at 96 kHz and 200 kHz, Hass 2021 (unpublished data) found individual differences in the upper frequency bands of the chirp above 15 kHz. As our on-collar recorders sampled up to 24 kHz, it is possible that we did not capture these higher frequency differences. Previous studies have shown that coatis maintain differentiated social relationships, with some individuals receiving disproportionate aggression. Given that coatis typically live in dense, forested environments, recognising individuals by their vocal signature likely plays a key role both in avoiding aggressive individuals and providing coalitionary support to individuals that cannot be clearly seen (Kaufmann 1962; Smith 1977; Gompper 1997; Gompper et al. 1997).

Within the Galaxy group, we observed significant variation in calling rates both within and between age-sex classes. Overall, contact calls were emitted at high rates for all group members, suggesting that all group members play a role in producing calls that help maintain group cohesion. Among the adult females of the group, there was high variation in contact call rate: two adult females emitted *chirp grunts* around three times more frequently than the other adult females, whereas these other adult females emitted *chirps* at a higher rate compared to *chirp grunts* (Figure S10). Understanding the functions of the different components of contact calls will provide insight into how group members use vocalisations to influence and shape collective outcomes. Since these data were recorded during the dry season, which also coincides with the breeding season, the observed calling patterns may have been influenced by the male’s presence in the group. Future studies should examine how call rates vary across seasons, group compositions, and life history stages.

#### Future Directions

By integrating advanced data collection methods with unsupervised approaches, this study has presented a robust framework for defining a species’ vocal repertoire, laying the groundwork for future investigations in vocal communication in this species. To better understand how these vocalisations are used in social processes, future studies should empirically test the function of different call types through playback experiments in wild populations, varying both call types and call rate emissions. This will help determine how the structural and temporal aspects of vocalisations influence collective outcomes. Playback experiments can clarify whether individuals recognise conspecifics, providing insight into how animals navigate their social landscape. Furthermore, these experiments allow us to examine how collective decisions are made within groups, for example by assessing whether individuals who call most frequently exert greater influence, or whether the caller’s identity carries more weight in shaping group decisions. By teasing apart these vocal dynamics within groups, we can gain a deeper understanding of the evolution of communication strategies and how these are shaped by the social environment.

## Acknowledgements

We thank the Smithsonian Tropical Research Institute for permission to conduct the study in Panama. Thank you to Carolina Mitre Ramos, Brandol Ortega, and Lucía Torrez for helping with the capturing, collaring, and data collection on the coatis in Gamboa and Barro Colorado Island, Panama. Thank you to Rachel Page and Melissa Cano for their support during fieldwork in Panama. The Arizona study was coordinated through the Fort Huachuca Wildlife Office (USAIC & FH), under supervision of H. Sheridan Stone and veterinary staff. We thank Jason Roback, John Kaufmann, Mike Siedman, and Marty Tuegel for their field assistance in Arizona.

We thank Michael Sutor for providing us with audio recordings of cub vocalisations. Thank you to Patrick Paetzold for assisting us with the design of the 3D printed cases which housed the audio recorders. Thank you to Julia Plecher, Leonie Treyer, and Caroline Preuß for their assistance on labelling calls from the collar recordings. We thank Vlad Demartsev for his advice on structuring the initial manuscript draft. We also thank the members of the Communication and Collective Movement group as well as members of the Communication and Coordination Across Scales team for their valuable feedback. Finally, we thank members of the department for the Ecology of Animal Societies for useful discussion.

