## Supplementary materials for "The vocal repertoire of white-nosed coatis: structural features, temporal dynamics, and social variations"

Appendices

Spectrogram generation

Spectrograms were generated using an FFT window length of 0.03 seconds and a hop length of 0.00375 seconds. The 'hanning' window function was applied to each audio frame (30 ms duration). When generating the mel filter bank, the frequency range up to half the minimum sample rate (the Nyquist frequency) was used. Spectrogram frequencies were binned into 40 logarithmically spaced bins (the “mels”) for frequency representation. The spectrograms were then denoised via median subtraction. To obtain numeric vectors for the UMAP embedding, the spectrograms were z-transformed, then zero-padded (so each call had equal length), and then concatenated. We used two dimensions for latent space representation.

Table S1. Details of audio data collection across study groups and locations

| **Study group** | **Location** | **No. individuals recorded** | **Date** | **Recording period** | **Recording device** | **hours labelled** | **Sample Rate (Hz)** | **No. calls used in acoustic features analysis** |
| --- | --- | --- | --- | --- | --- | --- | --- | --- |
| **Smarties** | BCI | 2 | 21.02.20 – 09.03.20 | 06:00 – 10:00 | Collar - TS Market, Edic Mini Tiny+ A77 | 48 | 22050 | 173 |
| **Galaxy** | SNP | 11 | 23.12.21 – 10.01.22 | 06:00 – 09:00 | Collar - Soroka 18E | 68 | 24000 | 178 |
| **Trago** | SNP | 9 | 22.03.22 | 16:00 – 17:00 | Manual -Ultrasound recorder | >1 | 250000 | 55 |
| **Galaxy** | SNP, when in trap | 5 | 21.02.22 | 22:00 – 23:00 | Manual - Phone mic | >1 | 48000 | 124 |
| **Troop 1** | AZ | >16 | 29.10.94 | 13:00 – 14:00 | Manual - Video camera | 16 min | 32000 | 26 |
| **Troop 3** | AZ | 11 | 30.07.96 | NC | Manual - Video camera | 3 min | 32000 | 25 |
| **Troop 5** | AZ | 6 | 01.07.99 | NC | Manual | 65 min | 32000 | 15 |
| **Troop 7** | AZ | 1 | 01.05.97 | NC | Manual - Video camera | 8 sec | 32000 | 11 |

Audio data were collected from wild white-nosed coati groups in Panama (Soberania National Park, SNP; Barro Colorado Island, BCI) using audio loggers, and in Arizona, USA (AZ) using manual recording devices. Note that 'NC' indicates that the information was not collected.

Table S2. Number of calls used for the original space and latent space representation of vocalisations

| **Label** | **Call count** |
| --- | --- |
| **Chirp** | 19361 |
| **Chirp grunt** | 10616 |
| **Chitter** | 4537 |
| **Chirp click** | 2488 |
| **Grunt** | 1506 |
| **Pew** | 1251 |
| **Squeal** | 354 |
| **Bop** | 293 |
| **Bark** | 166 |
| **Click** | 104 |
| **Hum** | 91 |
| Growl | 22 |
| Vibrate | 18 |
| Snort | 9 |

These calls were manually labelled from the collar recordings from 11 members of one group of white-nosed coatis (Galaxy group) in Soberania National Park, Panama. Labels in bold were included in the analysis.

Table S3. Call features of the 15 call types of the white-nosed coati repertoire

| **Call type** | **Previously described** | **Call count** | **Recording type** | **Location** | **Duration** | **Dominant frequency** | **IQR** | **Spectral flatness measure** |
| --- | --- | --- | --- | --- | --- | --- | --- | --- |
| chirp | Y | 34 | M | P, A | 0.087 ± 0.016 | 12474.494 ± 4490.659 | 10299.517 ± 4183.821 | 0.176 ± 0.138 |
| click | Y | 52 | L | P | 0.026 ± 0.003 | 1700.599 ± 1408.965 | 3168.592 ± 832.773 | 0.45 ± 0.068 |
| grunt | Y | 46 | L, M | P | 0.05 ± 0.013 | 250.361 ± 102.98 | 2467.663 ± 1443.221 | 0.342 ± 0.137 |
| chitter | Y | 53 | M | P, A | 0.063 ± 0.05 | 6540.242 ± 2180.585 | 8028.744 ± 4579.238 | 0.147 ± 0.092 |
| squeal | Y | 41 | L | P | 0.066 ± 0.037 | 2012.567 ± 1812.887 | 2915.911 ± 1066.893 | 0.521 ± 0.143 |
| growl | Y | 33 | L | P | 0.266 ± 0.122 | 228.382 ± 62.302 | 3531.445 ± 1583.715 | 0.523 ± 0.17 |
| bark | Y | 93 | L, M | P, A | 0.109 ± 0.022 | 4252.148 ± 1830.051 | 5489.085 ± 906.408 | 0.235 ± 0.25 |
| pew | N | 76 | L | P | 0.039 ± 0.006 | 2909.951 ± 2142.466 | 4019.531 ± 1358.569 | 0.434 ± 0.116 |
| hum | N | 15 | L | P | 0.131 ± 0.085 | 324.434 ± 128.032 | 3235.723 ± 1619.342 | 0.476 ± 0.11 |
| bop | N | 31 | L | P | 0.046 ± 0.012 | 214.718 ± 266.755 | 5618.952 ± 1401.541 | 0.594 ± 0.122 |
| vibrate | N | 35 | L | P | 0.126 ± 0.115 | 601.699 ± 309.41 | 3896.895 ± 1628.65 | 0.57 ± 0.14 |
| chop-chop | Y | 50 | M | P | 0.04 ± 0.011 | 837.879 ± 1652.697 | 3298.395 ± 1244.529 | 0.176 ± 0.087 |
| roar | Y | 6 | M | P, A | 0.341 ± 0.156 | 646.484 ± 472.625 | 1960.059 ± 1217.493 | 0.093 ± 0.071 |
| snort | Y | 31 | M | P, A | 0.152 ± 0.068 | 404.561 ± 448.456 | 3045.628 ± 1808.47 | 0.139 ± 0.151 |
| squeak | Y | 11 | M | A | 0.04 ± 0.006 | 2727.273 ± 1744.08 | 3673.295 ± 280.681 | 0.019 ± 0.003 |

For recorder type, L represents audio loggers attached to collars, M represents manual recordings from hand-held microphones. The location column defines where the call recordings were collected from, P stands for Panama, and A stands for Arizona. The IQR (interquartile range) column provide information for the frequency distribution of each call.

Table S4. Call types emitted in multi syllable calls sequences, and call types emitted in fast succession repetitively

| **Complex call name** | **complex call type** | **context** |
| --- | --- | --- |
| chirp-grunt | multi syllable | contact call |
| chirp-click-grunt | multi syllable | contact call |
| chirp-click-grunt-snort | multi syllable | contact call |
| chirp-click | multi syllable | contact call |
| bark (sometimes preceded by chirps) | Repeated | alarm |
| snort | Repeated | alarm |
| chop-chop | Repeated | alarm |
| chitter | Repeated | Aggression or distress |
| squeal | Repeated | aggression |
| pew | Repeated | excitement |

Table S5. Call types with associated call contexts

| **Call type** | **Context** |
| --- | --- |
| Chirp | Contact |
| Click | Contact |
| Grunt | Contact |
| Squeak | Excitement |
| Pew | Excitement |
| Chitter | Aggression/Distress |
| Squeal | Aggression |
| Growl | Aggression |
| Roar | Aggression/Alarm |
| Chop-chop | Conflict |
| Bark | Alarm |
| Snort | Alarm |
| Bop | Unknown |
| Hum | Unknown |
| Vibrate | Unknown |

Some calls are used in various contexts, but the contexts described here represent the most common situations in which these calls are used.

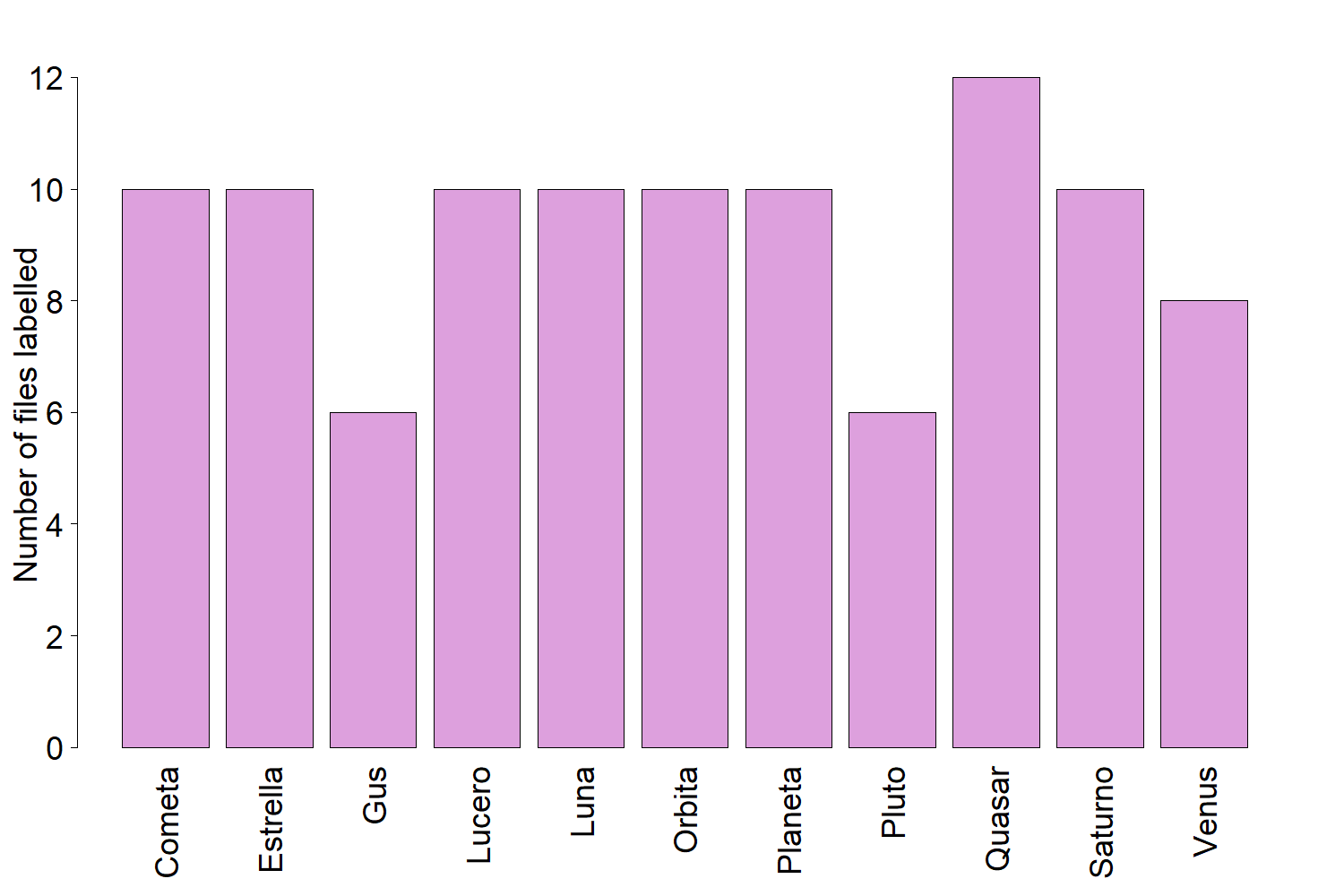

Figure S1. Number of files manually labelled from the audio logger recorders for each member of the Galaxy group. Each file had the last hour labelled, from 0800 to 0900 h.

(a) (b)

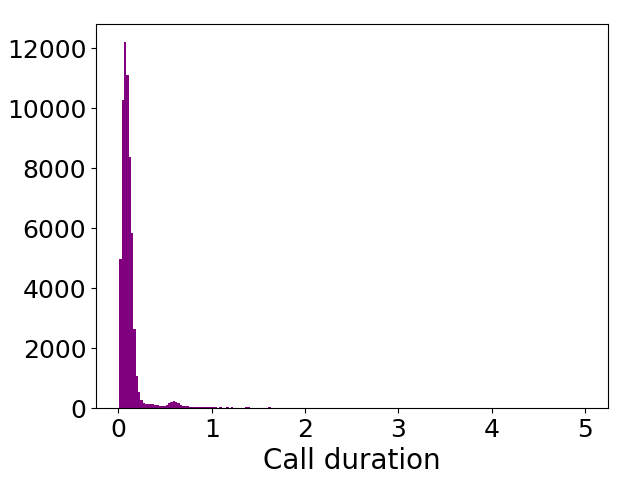

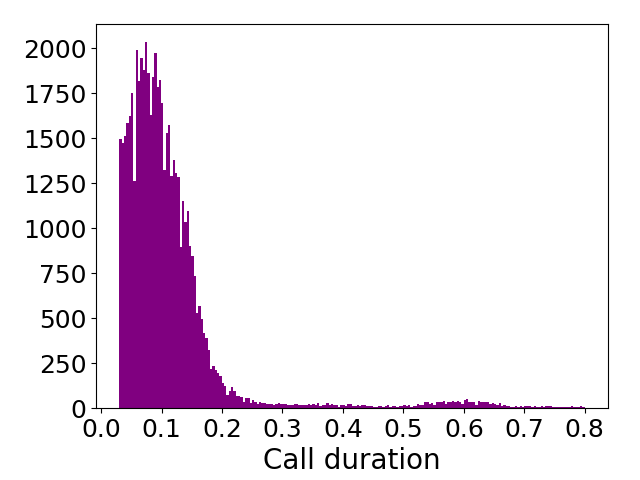

Figure S2. Histogram of call durations from the audio logger recordings used for UMAP and original spectrogram analysis before (a) and after (b) filtering calls to durations between 0.03 and 0.8s.

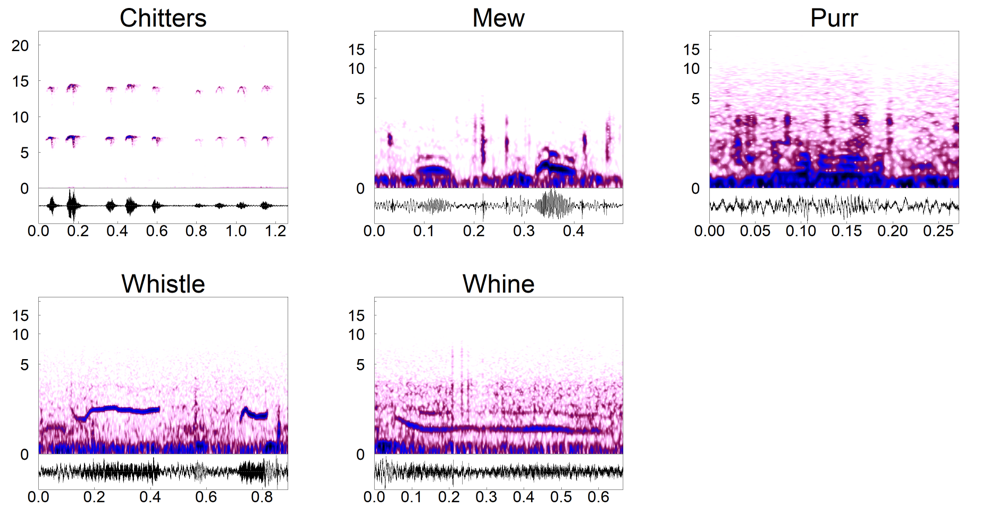

Figure S3. Spectrograms and corresponding waveforms of wild nestling calls recorded from a manual recorder by Michael Sutor when cubs were nursing in the nest, Arizona. Nestlings were 2 – 4 weeks old (Sutor 2000).

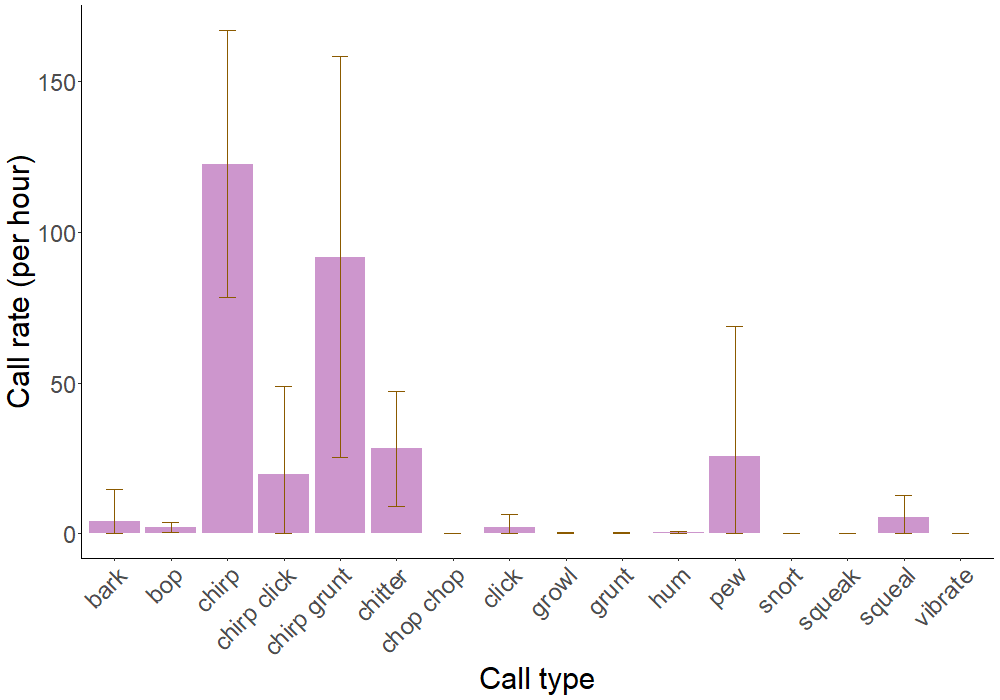

Figure S4. Mean call rates for the most commonly emitted multi syllable calls and single syllable calls for all group members in the Galaxy group. Calls were recorded on audio logger recorders and manually labelled. Error bars represent the standard deviation of mean call rates of all group members. Rare call types such as the *roar* were not found in these audio logger recordings and therefore were not included. Calls were recorded during the mating season and may not reflect call rates for other times of the year.

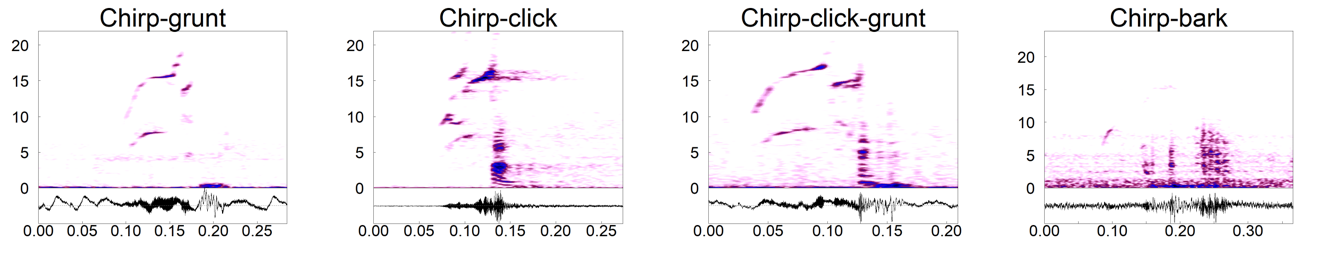

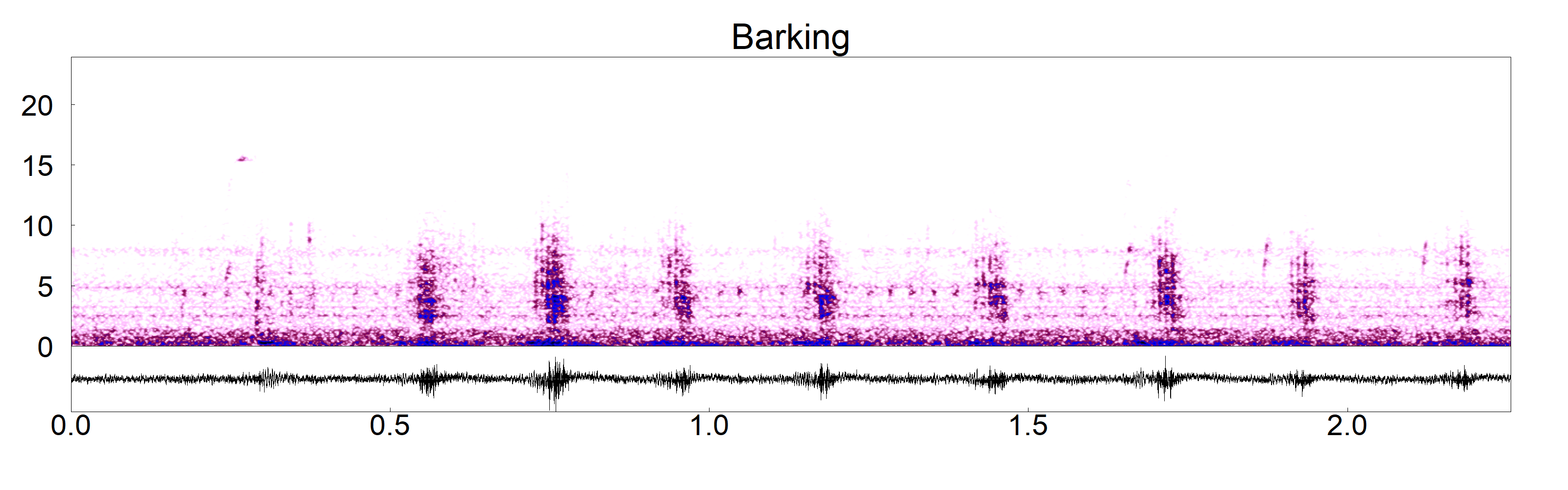

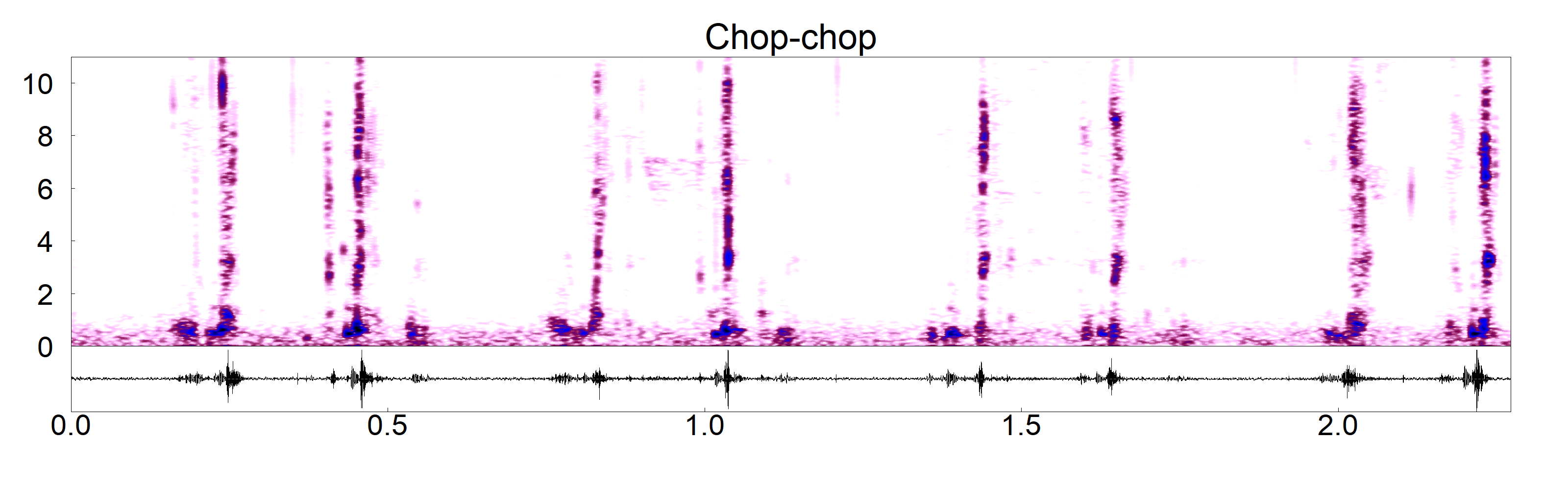

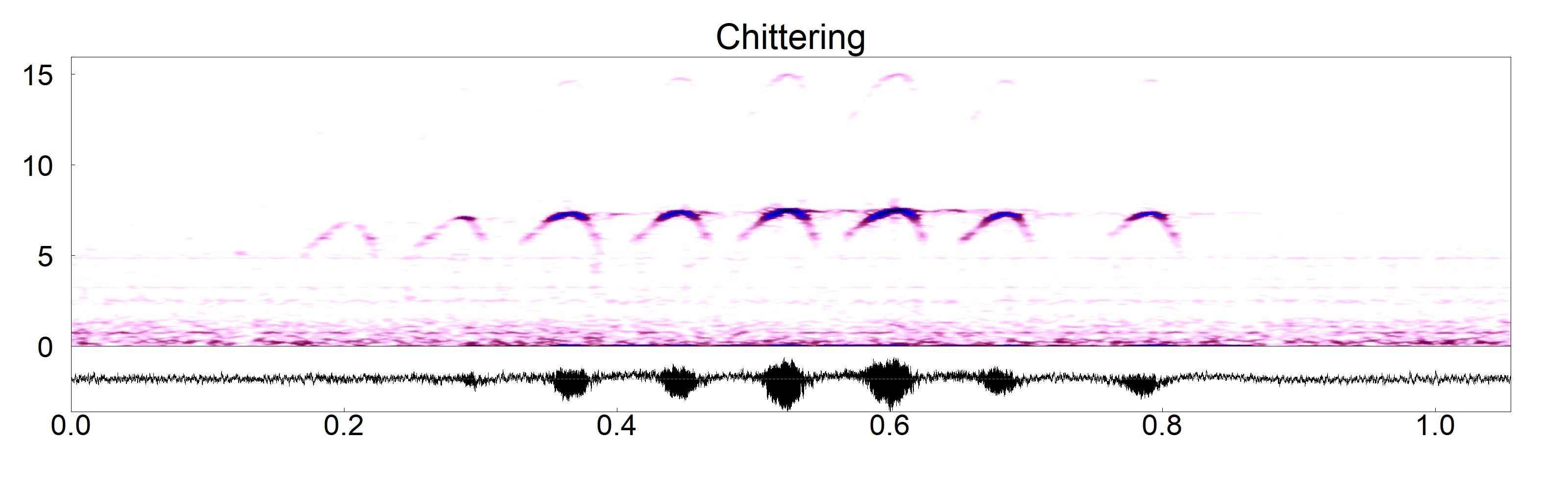

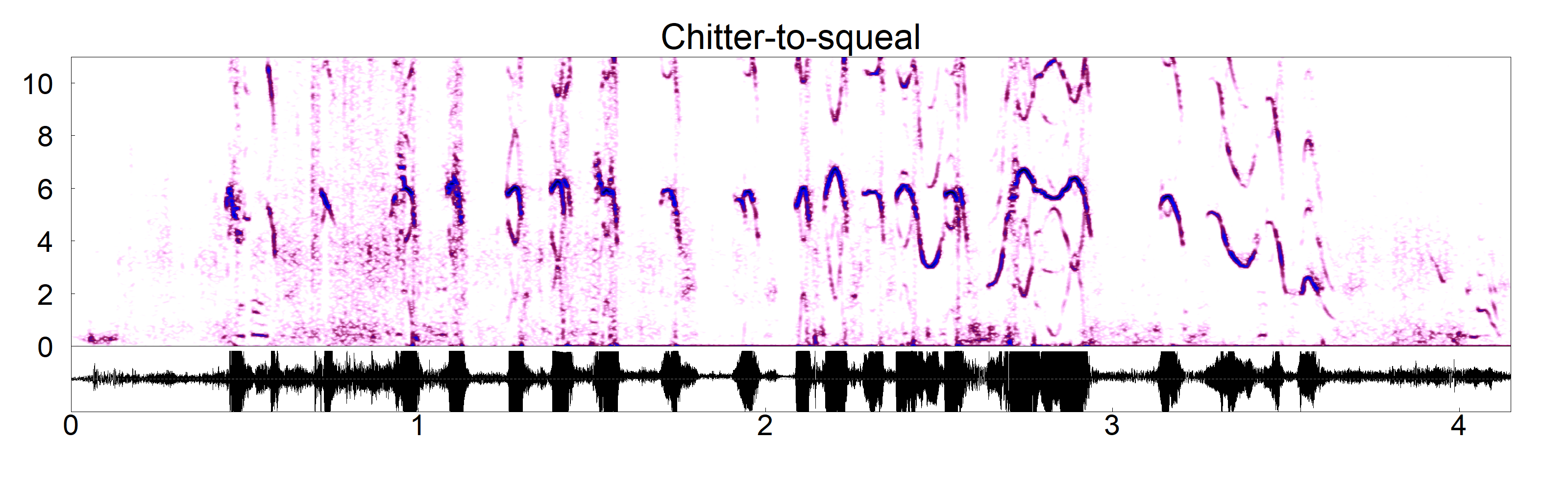

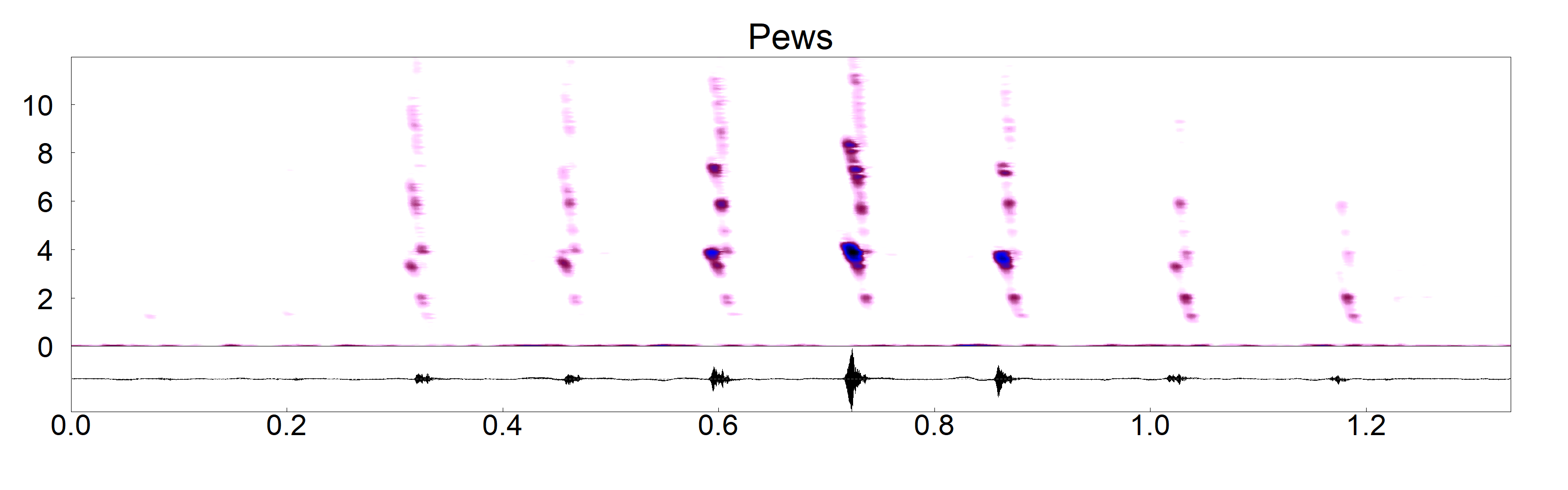

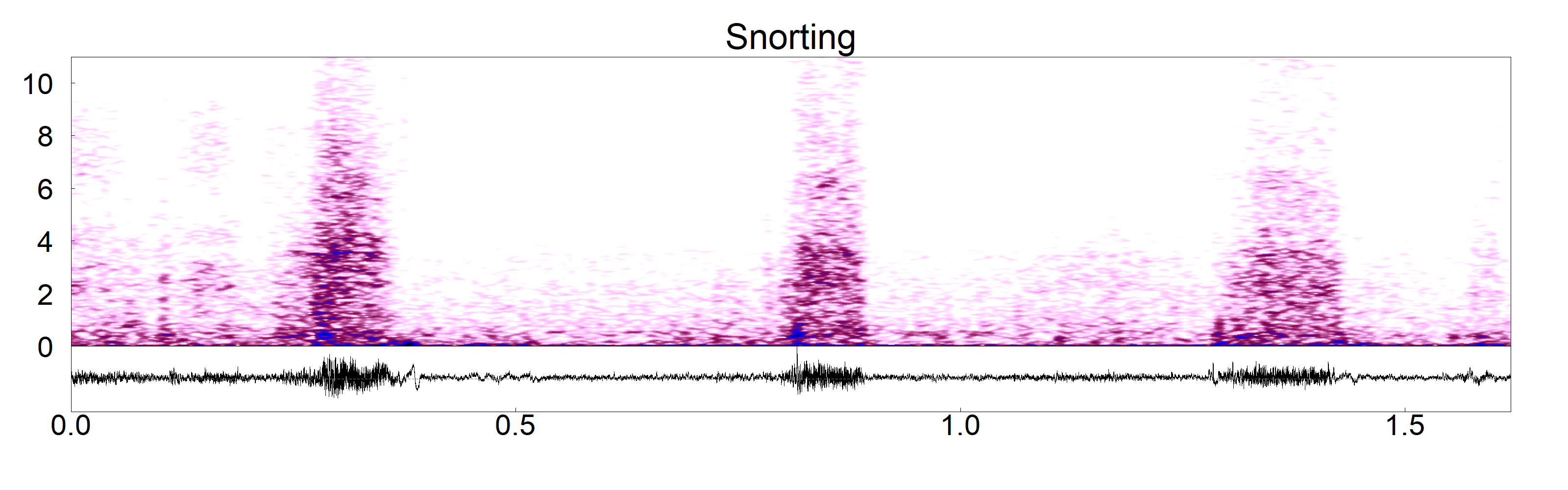

Figure S5. Spectrograms and corresponding waveforms of multi-syllable calls and calls emitted repetitively in fast succession, the x-axis represents time in seconds, and the y-axis indicates frequency in kHz.

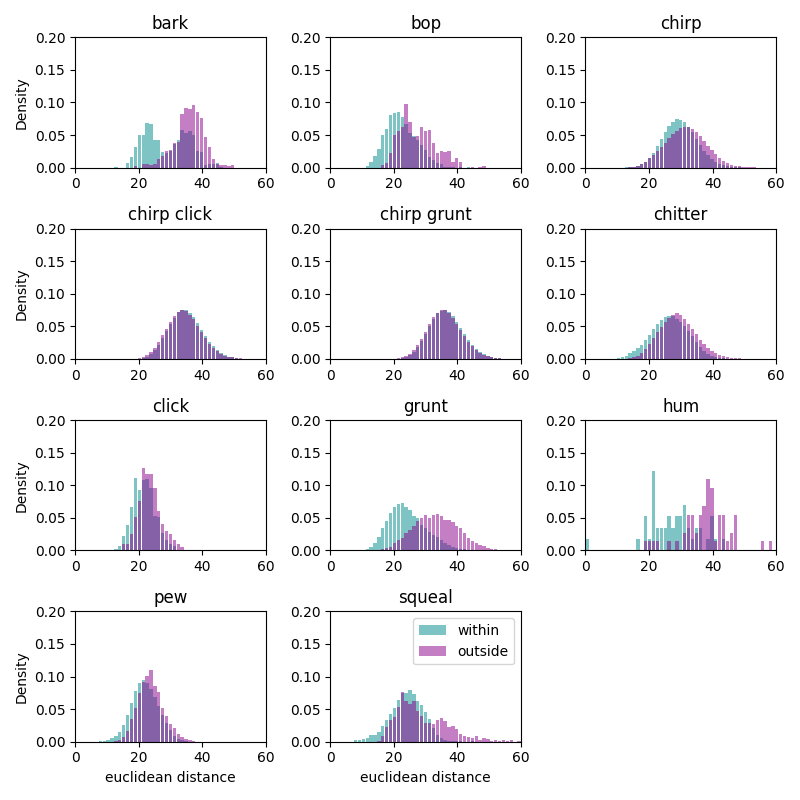

Figure S6. The distribution of pairwise distances in the original space within each call type compared to different call types.

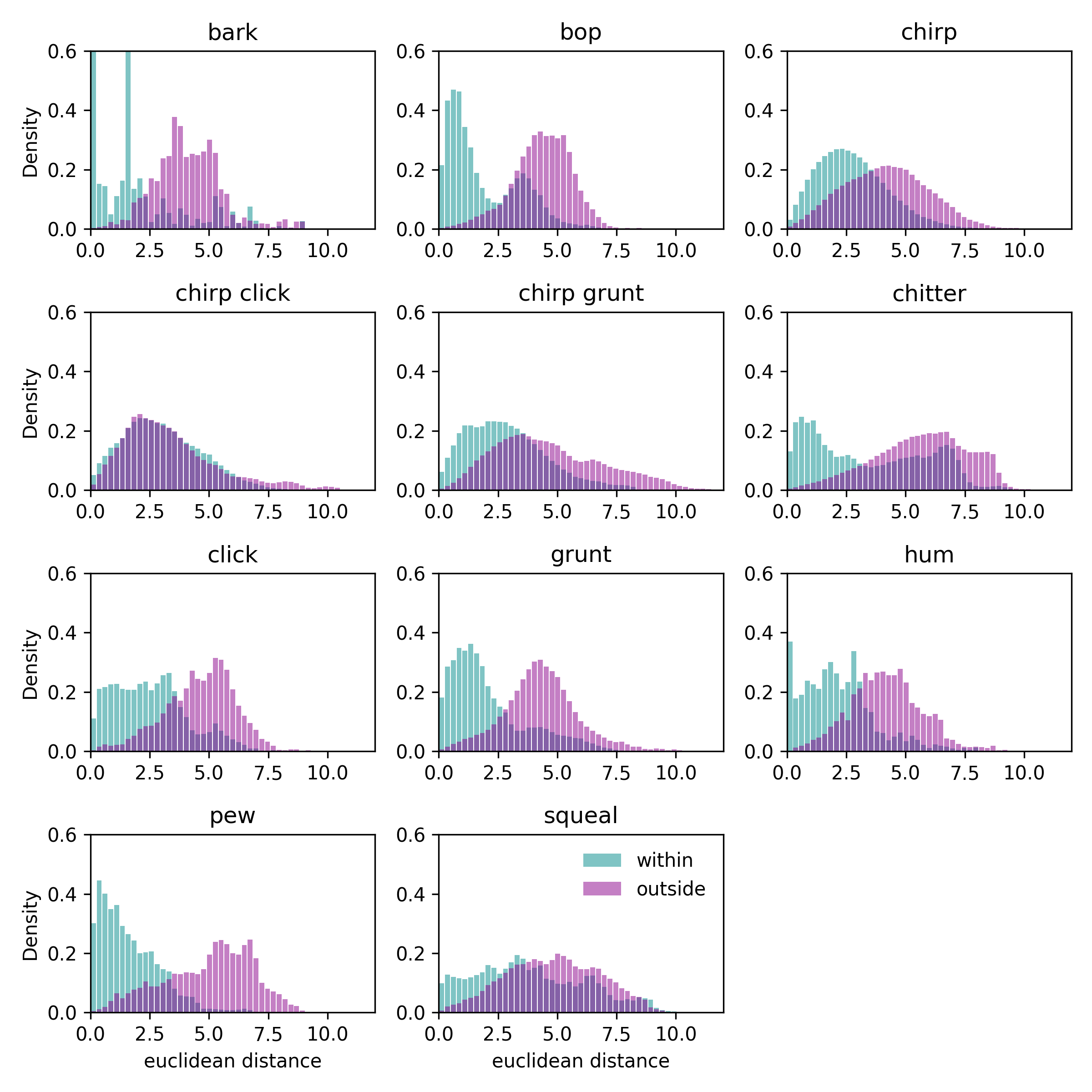

Figure S7. The distribution of pairwise distances in the 2D UMAP space within each call type compared to different call types.

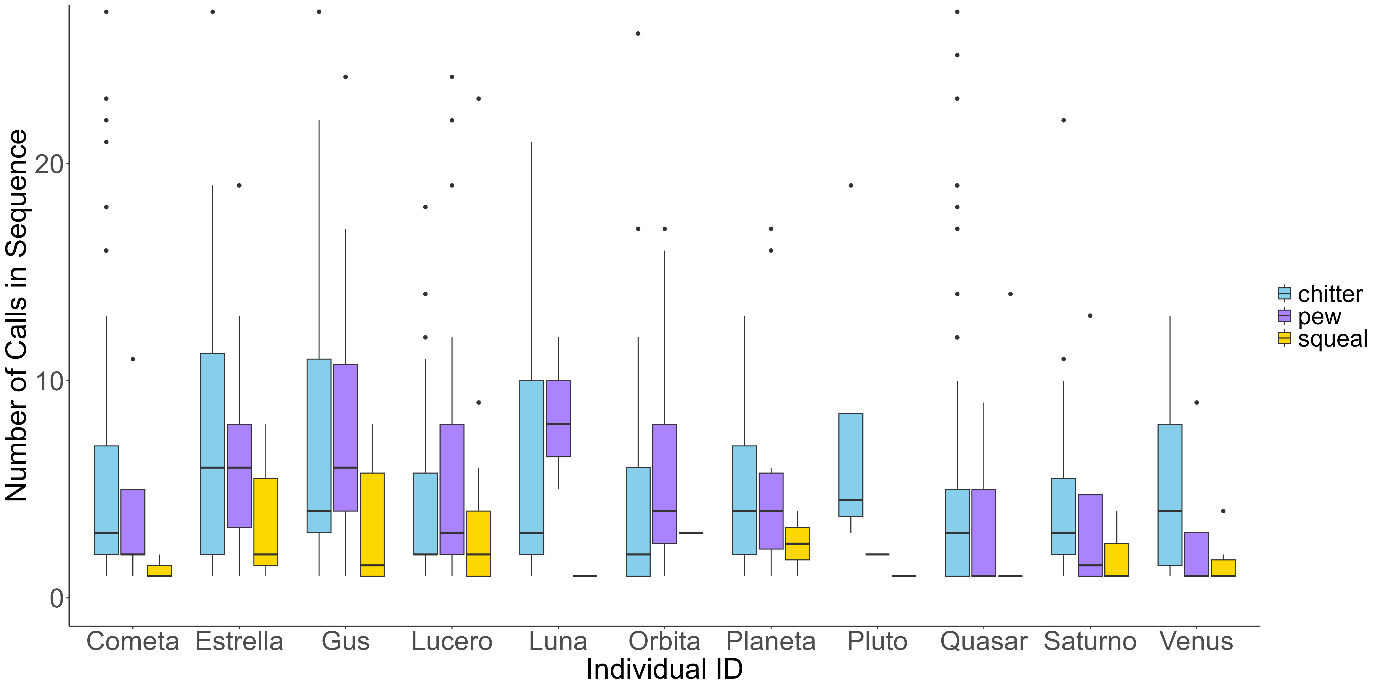

Figure S8. Distribution of the number of calls per sequence for three call types—*chitter, pew*, and *squeal*—among group members in a single group of white-nosed coatis (Galaxy group) in Soberania National Park, Panama.

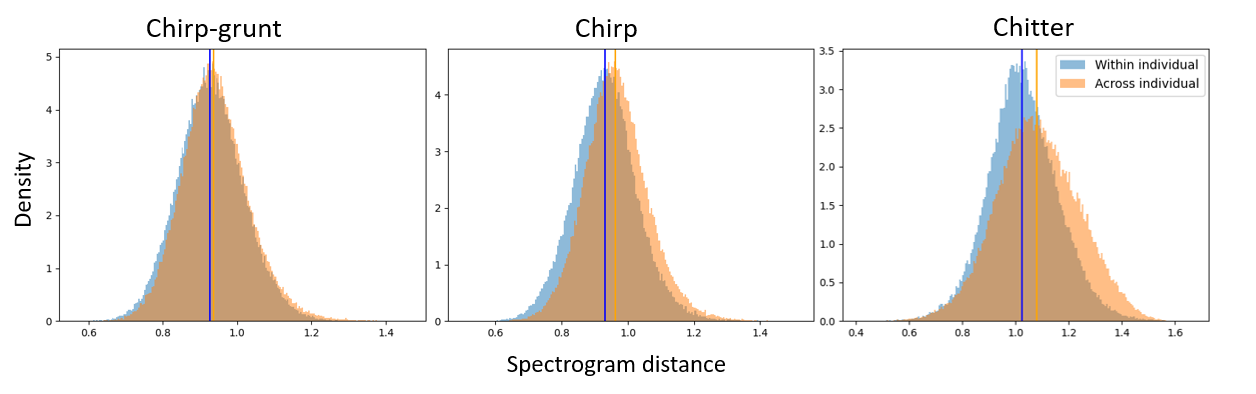

Figure S9. Distributions of pairwise spectrogram distances for the most common call labels, comparing calls recorded from the same individual on different recorders (within individual) to calls recorded from different individuals on different recorders (across individual). Distances were calculated from vocalisations of all members of a single group of white-nosed coatis (Galaxy group) in Soberania National Park, Panama.

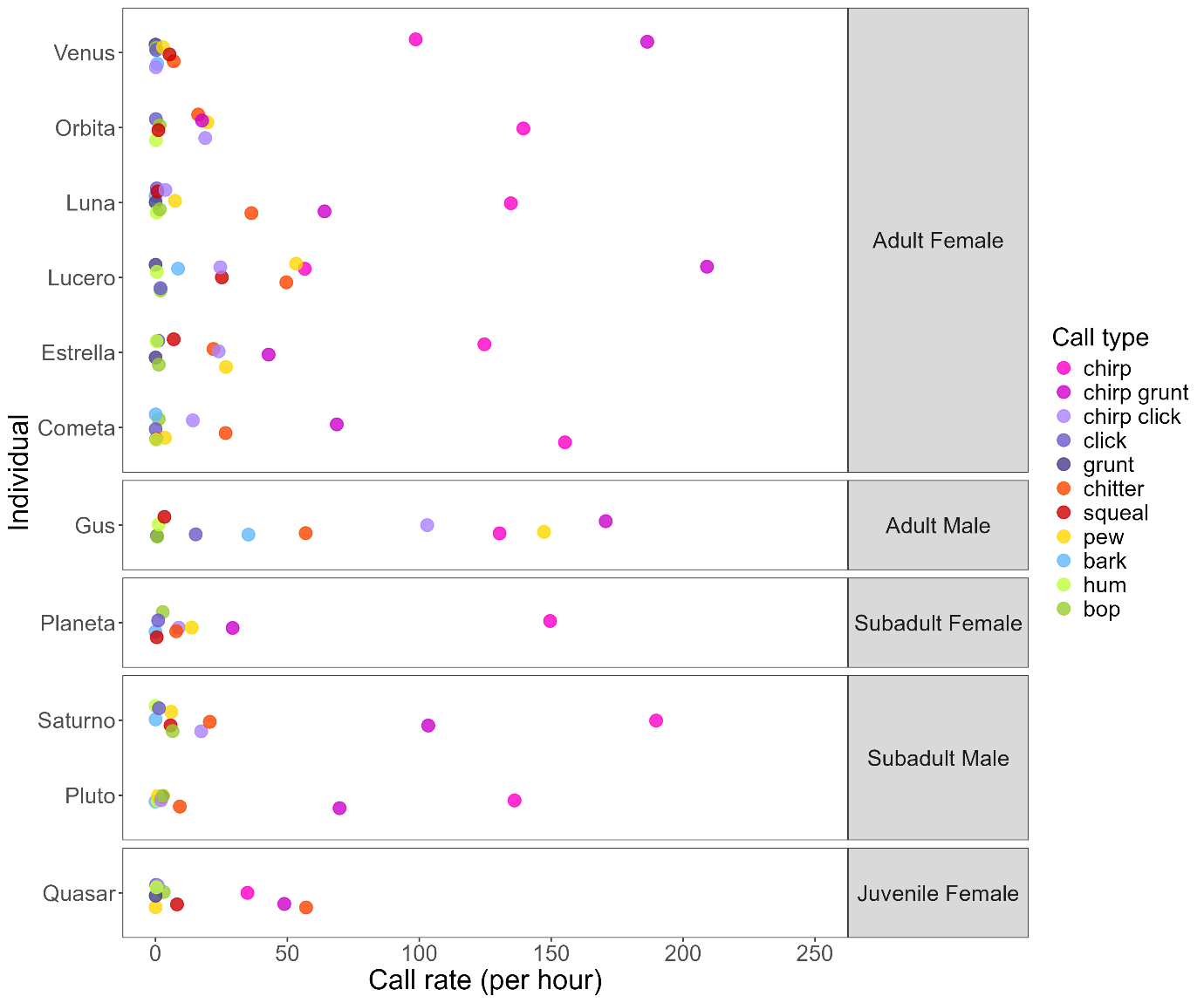

Figure S10. Mean calling rates of all 11 members of the Galaxy group in Soberania National Park, Panama.
